# Iterative engineering of a compact Cas9 ortholog for *in vivo* gene editing via single AAV delivery

**DOI:** 10.64898/2026.01.12.698990

**Authors:** Alessandro Barenghi, Jamie A. Hackett, Neil Humphreys, James A. Sawitzke

## Abstract

The development of compact and efficient CRISPR-Cas systems is crucial for biomedical and therapeutic genome editing, particularly *in vivo* applications based on viral delivery. Here, we performed a comparative functional screen of seven Cas9 orthologs to systematically evaluate their genome editing activity in mammalian cells. Among these, Cme2—a 1008 amino acid nuclease recognizing a 5′-NAGNGC PAM—emerged as a promising candidate based on its compact size and baseline editing activity. To overcome its limited native efficiency, we employed a dual engineering approach combining sgRNA scaffold optimization and rational protein mutagenesis. The resulting variant, *en*Cme2, exhibits markedly improved editing efficiency across multiple loci in both mouse and human cells while maintaining extremely high specificity and minimal off-target activity. Importantly, the small size of *en*Cme2 permits packaging of the complete system into a single rAAV vector, enabling efficient genome editing/HDR in *in vivo* tissues and mouse embryos, and facile generation of transgenic models. These results establish *en*Cme2 as a compact, precise, and AAV-compatible genome editing platform with broad applicability for *in vivo* research and therapeutic approaches, especially where high specificity is desirable.

## INTRODUCTION

The emergence of CRISPR (Clustered Regularly Interspaced Short Palindromic Repeats) technology has transformed genome editing by enabling programmable, sequence-specific manipulation of DNA across virtually all biological systems [1, 2]. At its core, CRISPR–Cas systems rely on RNA-guided nucleases to achieve sequence-specific targeting of genomic DNA, enabling precise induction of double-strand breaks (DSBs). In type II CRISPR-Cas9 systems, target recognition is naturally mediated by a dual-RNA complex composed of the CRISPR RNA (crRNA), which guides Cas9 to the target sequence, and the trans-activating crRNA (tracrRNA), which helps form the functional RNA-Cas9 complex. For genome engineering applications, these two RNAs are typically combined into a single-guide RNA (sgRNA), which integrates target recognition and Cas9 scaffolding into a single molecule [2]. Following a DSB, repair of the lesion typically proceeds via non-homologous end-joining (NHEJ), introducing stochastic insertions and deletions that can disrupt coding sequences, or via homology-directed repair (HDR), which permits precise sequence replacement or insertion [3]. More recent CRISPR systems employ Cas9 nickase (nCas9) or catalytically inactive Cas9 (dCas9) fused to effector enzymes. For example, base editors couple nCas9 to cytidine or adenine deaminases to catalyse single-nucleotide conversions without introducing DSBs, whereas prime editors utilise a reverse transcriptase and prime-editing guide RNA (pegRNA) to directly write new genetic information into the genome. Collectively, these CRISPR-derived technologies have had a transformative impact across basic research, agriculture and medicine, with therapeutic applications now advancing into clinical development and patients [4–8].

While CRISPR has opened unprecedented opportunities, the efficiency and accuracy of genome editing are influenced by the choice of delivery method. Adeno-associated viruses (AAVs) are widely used for genome editing because they combine efficient delivery with safety and versatility. Their natural and engineered serotypes provide both broad and tissue-specific tropisms, making them adaptable for both gene therapy applications and the generation of transgenic animals [9–12]. For the latter, recombinant AAV (rAAV) delivery is minimally invasive, requires no specialised equipment, improves embryo survival, and improves the efficacy of large-scale genomic modifications [13]. However, AAV’s limited packaging capacity (<4.7 kb) presents a significant hurdle for delivering the full repertoire of CRISPR components based around *Streptococcus pyogenes* Cas9 (SpCas9; ∼4.1 kb), particularly for applications requiring multiple sgRNAs, HDR templates or larger precision tools such as base, prime or epigenome editors. To overcome the inherent challenges of CRISPR delivery, efforts have focused on developing smaller Cas9 variants such as CjCas9 [14], Nme2Cas9 [15], SaCas9 [16] and SauriCas9 [17], Cas12 family members [18, 19] and transposon-encoded TnpB proteins [20, 21]. Despite the successful *in vivo* application of these systems via AAV delivery, they present limitations relative to conventional SpCas9, including low editing efficiency, restrictive PAM requirements and highly variable gRNA functionality. Moreover, the development of highly precise systems is crucial to minimizing the risk of off-target effects. To overcome these functional and delivery limitations, the ongoing development of novel and compact CRISPR tools with enhanced features is essential.

In this study, we evaluate the editing efficiency of a panel of recently identified compact Cas9 orthologs, whose *in vivo* functionality remains largely unexplored [22]. Among them, Cme2 (∼3 kb; 1008 aa), which recognizes the 5’ NAGNGC PAM sequence, exhibited the highest activity and was selected for further optimization. Through extensive sgRNA and protein engineering, we developed an optimized variant, *en*Cme2, which displays enhanced editing efficiency, an expanded targeting range, and tremendous on-target specificity. By leveraging its compact size for single all-in-one rAAV delivery, we demonstrate that *en*Cme2 enables efficient genome editing, including HDR, across endogenous targets and in mouse embryos. Our findings establish *en*Cme2 as a versatile and highly specific genome-editing tool, offering a compact alternative to existing CRISPR systems.

## RESULTS

### Systematic characterization of a panel of compact Cas9 orthologs

To identify promising candidates for miniaturized CRISPR systems, we first sought to examine unrelated Cas9 orthologs from different organisms with evidence of *in vitro* activity [22]. These systems were primarily selected based on their compact size (∼1000 amino acids; 3 kb) and a relatively relaxed PAM sequence structure (Figure 1A-B). Importantly, while the activity of each enzyme has been implicated using purified components *in vitro* [22], the extent to which they function at genomic targets in living cells is unknown. To test this, we first employed a mouse embryonic stem cell (mESC) line wherein the *trp53* gene carries an in-frame knock-in of tdTomato, enabling endogenous expression to be tracked [23]. We systematically engineered each Cas9 system under a dox-inducible promoter (TRE3G) to enable temporal control over editing activity, and then genomically integrated them into *p53*^tdTomato^ mESC using the piggyBac system. Specifically, for each line we introduced one Cas9 ortholog, in tandem with one of three ortholog-specific sgRNA that target the *tdTomato* sequence (Figure 1C). Given limited prior knowledge, sgRNAs were selected based on optimal PAM sequences, without taking into account other factors such as target sequence or the genomic context, to remove potential bias [24].

**Figure 1.**
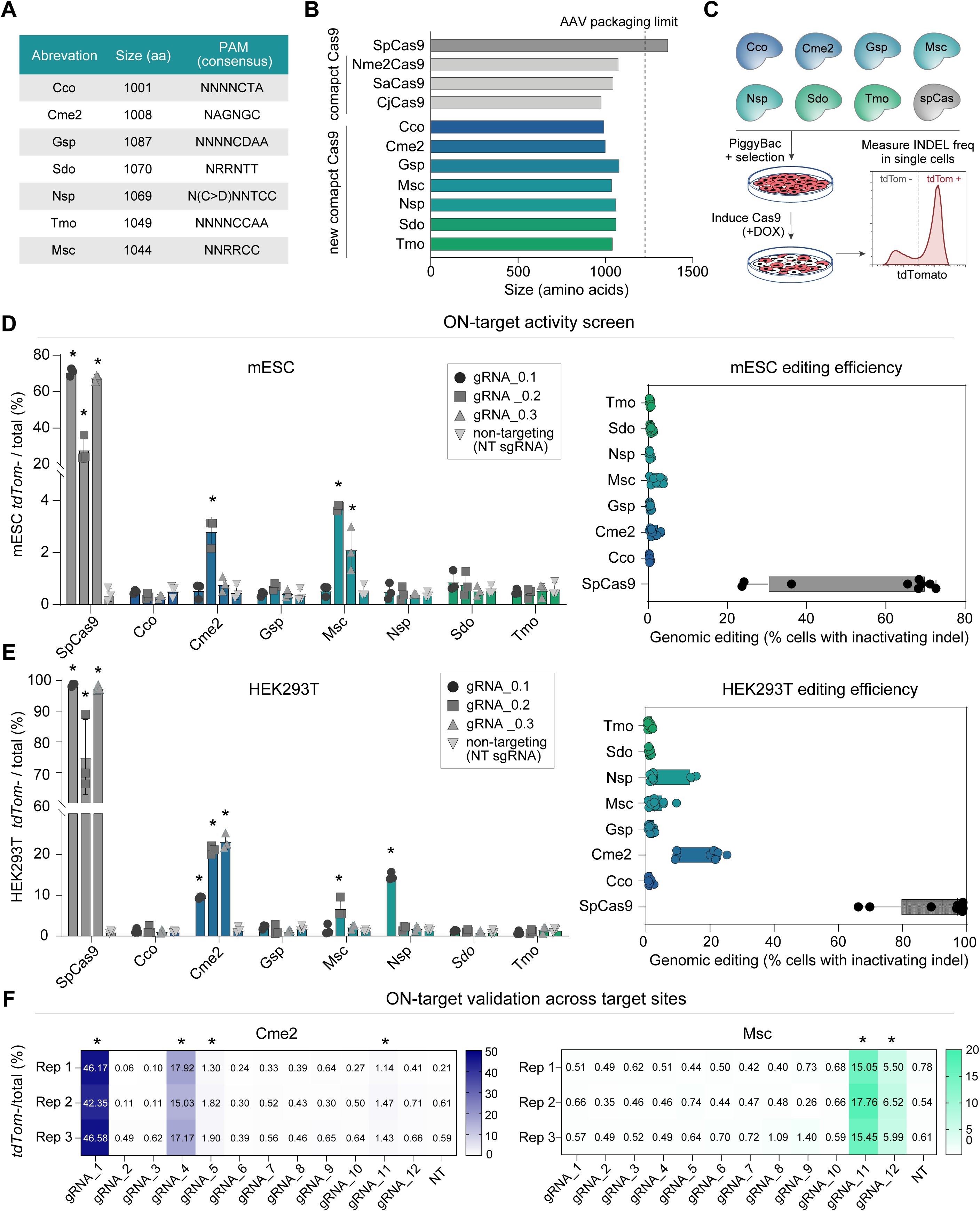
Screening compact Cas9 for editing efficacy. (A) Table of the Cas9 orthologs utilized in this work, showing their size (amino acid) and PAM sequence (5’). (B) Schematic size comparisons of the panel of Cas9 orthologs tested with SpCas9 and other compact CRISPR systems commonly used for *in vivo* genome editing. (C) Schematic outlining the experimental steps to evaluate Cas9 ortholog editing efficiency by *tdTomato* knock-out (KO), quantified by flow cytometry analysis. PiggyBac constructs encode for: ortholog-specific hU6-sgRNA cassettes and puromycin selection gene; doxycycline-inducible Cas9 orthologs and hygromycin selection gene. (D–E) Comparison of SpCas9 cleavage efficiency with Cco, Cme2, Gsp, Msc, Nsp, Sdo and Tmo Cas9 orthologs using three ortholog-specific spacer sequences targeting the *tdTomato* gene in mESC (D) and HEK293T cells (E). Editing efficiencies at individual target sites are shown in the left panels, while mean editing efficiencies for each Cas9 ortholog across the three sites are presented in the right panels. (F) Heatmaps showing editing efficiencies of Cme2 (left) and Msc (right) using twelve distinct sgRNAs, quantified by flow cytometry. Asterisks indicate P-values from unpaired *t*-test comparing three independent biological replicates to the respective controls. *P < 0.05. Error bars ± SD.

We first determined the editing efficiency of each Cas9 ortholog by quantifying the fraction of single cells that become tdTomato-negative, using flow cytometry. This strategy is facile and evaluates functionally-relevant *de novo* DNA insertions/deletions (indels), albeit underestimates Cas9 editing activity due to in-frame mutations that retain fluorescence. After 3 days of DOX-induction, we observed moderate ON-target editing efficiency for two of the seven tested Cas9 orthologs (Cme2 and Msc) with at least one sgRNA, whilst other systems exhibited only background activity (Figure 1D). SpCas9 outperforms all orthologs, with a peak editing of 72.6% (mean=55.4% ± 20.9%), compared to 3.1% for Cme2 (mean=1.4% ± 1.1%) and 3.8% for Msc (mean=2.13% ± 1.45%). We confirmed that all Cas9 proteins are well expressed and stable in mESC, with any difference in protein levels not correlated with enzymatic activity, suggesting the absence of activity reflects sub-optimal enzymatic kinetics rather than destabilized Cas9 proteins (Supplementary Figure 1A). To investigate the panel of Cas9 orthologs within different cell types/species, we next generated a transgenic human HEK293T cell line in which *tdTomato* was inserted into the AAVS1 safe harbor locus. Using the same sgRNAs, we observed a maximum editing efficacy of 22.5% for Cme2 (mean=17.9% ± 6.1%) and 9.58 for Msc (mean=3.5% ± 2.6%), considerably higher than mESC. Notably, we observed that unlike mESC, Nsp appears to exhibit editing activity up to 15.7% in HEK293T cells (Figure 1E). These data imply that Cme2, Msc and to a lesser extent Nsp exhibit genomic editing activity at target loci in living cells, albeit at low levels. Moreover, they reveal their efficiency is not solely dependent on the sgRNA sequence but is also influenced by extrinsic factors [25, 26].

We therefore moved forward with characterization of Cme2 and Msc by first examining sgRNA variance with an additional twelve sgRNAs. This confirmed the functionality of both Cas9 orthologs in mESC, with 4/12 and 2/12 sgRNAs displaying significant activity (P < 0.05) for Cme and Msc, up to a maximum of 46.6% and 17.8% editing, respectively (Figure 1F). To investigate whether sgRNA expression levels could affect editing efficiency, we engineered *trp53^tdTomato^* mESC with increasing concentrations of sgRNA encoding plasmids. We observed that at least one of the three guides tested for each Cas9 ortholog resulted in a modest increase in editing efficiency (2.2-fold and 1.5-fold for Cme2 and Msc, respectively) at higher sgRNA concentrations (Supplementary Figure 1B-C). This suggests that the activity of the two Cas9 orthologs is primarily influenced by the choice of target and the specific protospacer sequence, albeit sgRNA levels/stability are partially limiting. Given its higher baseline activity and sgRNA tolerance, we decided to focus on the Cme2 system going forward.

### Enhancing Cme2 editing efficiency through gRNA engineering

We aimed to systematically optimize aspects of Cme2 protein function and sgRNA design architecture, with the goal of combining each iterative improvement to synergistically enhance overall activity. First, we focused on engineering the sgRNA by asking whether the spacer length influences Cme2 activity. We modified the length of Cme2 gRNA_1 and gRNA_4 spacers (hereafter referred to as gRNA_1 and gRNA_4) from 21 to 26 nucleotides (nt) at 1 nt resolution. Although the two sgRNAs exhibit different optimal lengths, 24nt provided the best balance between functionality and consistency (Supplementary Figure 2A), and was therefore selected as the preferred length for subsequent experiments. Notably, all sgRNAs exhibited no activity with spacer lengths shorter than 22 nt, in line with previous reports that Cas9 systems can require spacers longer than 20 nt for optimal function [14] [27–29].

Next, we used the RNAfold prediction tool from ViennaRNA to reconstruct the secondary structure of Cme2 sgRNA, aiming to identify potential modification sites [30]. We found Cme2 sgRNA consists of a repeat-antirepeat duplex, a nexus, and two 3’ tracrRNA hairpins (stem-loops 1 and 2) (Figure 2A and Supplementary Figure 2B). Using a structure-function guided approach, we modified each region of the sgRNA, starting from the repeat-antirepeat duplex. It has been reported that extension of the cRNA-tracrRNA duplex can increase editing efficiencies by facilitating gRNA-Cas9 assembly [31]. We therefore extended Cme2 sgRNA duplex up to 6 bp at a 1 bp resolution and scored the knockout (INDEL) efficiency using Cme2 gRNA_1 and gRNA_4 spacers (Supplementary Figure 2C). The data show a modest increase of editing when the duplex is extended by +1 bp in both sites tested, while further extension does not significantly affect functionality (Figure 2B). Thus, the Cme2_a1 variant was selected for inclusion in the final design.

**Figure 2.**
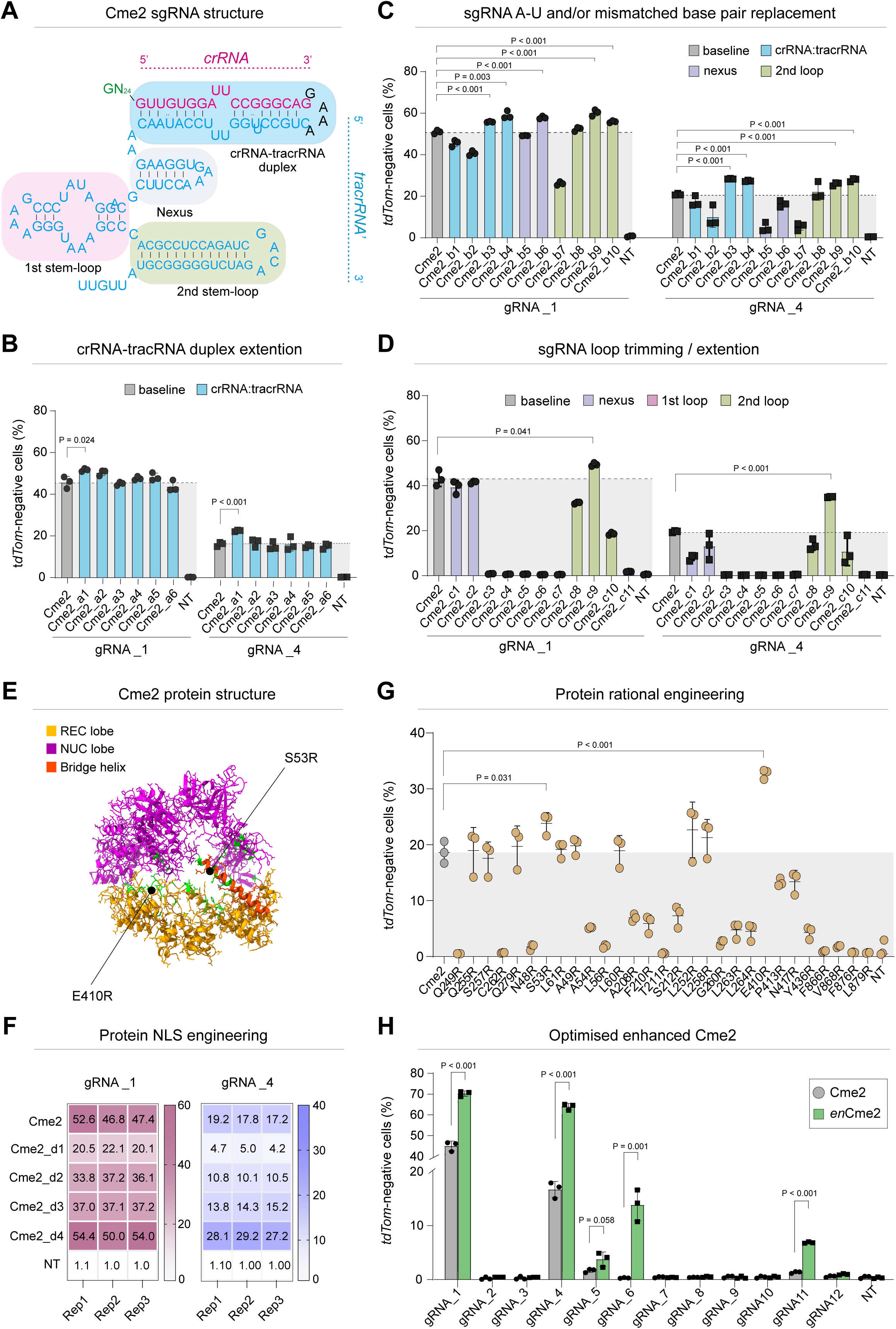
Systematic engineering of an enhanced *(en)*Cme2. (A) Schematic representing Cme2 sgRNA scaffold structure. Primary structures are highlighted as follows: blue (crRNA-tracrRNA duplex), grey (tracrRNA nexus), pink and green (3’ tracrRNA stem-loops 1 and 2, respectively). (B) Histogram showing the effects of crRNA-tracrRNA duplex extension (up to 6 bp) on Cme2 activity. (C) Histogram detailing the change of editing efficiency induced by A/U to C/G replacement and/or mismatch corrections throughout the Cme2 sgRNA scaffold. (D) Modification of the Cme2 sgRNA architecture through trimming/extension of the nexus region and the 3′ tracrRNA stem-loops 1 and 2 and their impact on Cme2 efficiency. (E) Predicted structure of Cme2 protein, color-coded as follows: the recognition lobe (REC lobe) is indicated in yellow, the nuclease lobe (NUC lobe) in purple and the bridge helix in red. Residues selected for mutagenesis are highlighted in green. The model was generated with AlphaFold v2.3.0 and visualized using ChimeraX 1.9. (F) Effects of nuclear localization signal (NLS) optimization on Cme2 editing. Three biological replicates are shown separately for each target sequence. (G) Editing activity of Cme2 protein variants carrying single arginine substitutions. (H) Histogram showing synergistic enhancement of editing activity by the combination of Cme2_a01, Cme2_b03, Cme2_b10 and Cme2_c09 sgRNA designs with E410R and Cme2_d04 protein variants (to obtain *en*Cme2) compared to Cme2 at twelve individual *tdTomato* target sites. In all panels, P-values were obtained using unpaired *t*-tests over three independent biological replicates; Error bars ± SD.

Next, we sought to increase sgRNA stability, since it has been proposed that sgRNA self-folding free energy and positional entropy can strongly influence cleavage activity [32]. Indeed, misfolded gRNAs are not only non-functional but also compete for Cas9 binding, thereby acting as a dominant-negative that reduces editing [33]. A strategy to ameliorate this is to replace mismatched base pairs in the sgRNA or correctly paired A-U matches with thermodynamically stable C-G [34, 35]. We identified ten potential sites throughout the whole sgRNA scaffold and subsequently modified their sequence (see Supplementary Figure 2D for details). Quantitative flow analysis revealed that the effects of each of these sgRNA scaffold sequence edits are consistent in both genomic sites tested, with some variants significantly enhancing editing. In particular, Cme2_b9 and Cme2_b10 variants, which target the 3′ tracrRNA stem-loop 2, exhibited average increases in editing efficiency of 21.2% and 21.6%, respectively. Similarly, Cme2_b3 and Cme2_b4 sgRNA variants enhanced editing activity by 15.2% and 16.8% on average, respectively (Figure 2C). Specifically, the latter two modifications were designed to correct a bulge in the crRNA-tracrRNA duplex, which led to a reduction of the positional entropy of the scaffold in the model (Supplementary Figure 2D).

Another strategy to enhance editing efficiency of CRISPR systems is to modify the length of the stem-loops formed by the tracrRNA as well as its 3’ end [36] [32] [19]. In this respect, the extension of a specific region could improve sgRNA interaction with the Cas9 protein, whilst the trimming could facilitate sgRNA folding and increase stability (for example, by removing structurally disordered regions in the sgRNA [19]). Taking into account the difficulty in predicting the outcome of the aforementioned strategy, we decided to broadly evaluate the effects of a partial and/or total deletion or extension of the tracrRNA stem regions. In particular, Cme2 tracrRNA nexus was extended or truncated by 2 bp, stem-loop 1 was trimmed by 3, 6 and 9bp, and stem-loop 2 was either extended by 3bp or truncated by 4 and 7bp (Supplementary Data). Moreover, the three stem regions were completely truncated. Most of the variants impaired editing efficiency, with complete truncation of any stem region nullifying Cme2 activity. However, a 4bp trimming of stem-loop 2 exhibited higher indel frequency in both sites tested (Figure 2D). Specifically, the Cme2_c9 variant led to a 1.2-fold increased editing for gRNA_1 and to a 1.8-fold increase in gRNA_4. Given the iterative improvement of each sgRNA engineering step, we tested their synergy by combining them to obtain the final sgRNA design. Cme2_a1, Cme2_b10 and Cme2_c9 variants were combined with either Cme2_b3 or Cme2_b4 to generate two different combinations, named respectively Cme2_en_sg1 and Cme2_en_sg2. Cme2_b9 was omitted from the final design because the Cme2_c9 variant abolishes the structural alteration introduced by Cme2_b9, while displaying a stronger effect. While we did not observe a synergistic relationship, both Cme2_en_sg1 and Cme2_en_sg2 exhibited a significantly higher level of editing relative to the starting sgRNA architecture and in line with the best level of any individual edit (Supplementary Figure 2E), suggesting an overall enhancement of Cme2 sgRNA scaffold structure.

### Enhancing *Cme2* editing efficiency through protein engineering

To complement optimization of sgRNA architecture, we sought to improve the Cme2 protein through guided mutagenesis and screening for variants with higher editing efficiency. The substitution of basic/neutral amino acids in the DNA or RNA recognition domains with positively charged residues (typically arginine, R) can improve editing by promoting protein-nucleic acid interaction and RNP formation [37–39]. In order to identify possible amino acids involved in Cme2 binding to nucleic acids, taking into account the unavailability of structural data, we generated a model of the protein using AlphaFold v2.3.0 (Figure 2E) [40]. The model was superimposed on the representation of other Cas9 orthologs of similar sizes whose structures have been resolved, including SaCas9, CjCas9 and Nme2Cas9 [41–43]. This, together with direct protein sequence alignment, allowed us to select twenty-nine amino acids in conserved regions having side chains internally facing Cme2 DNA binding pockets (Figure 2E). Next, we individually mutated those residues into arginine to obtain a mutant library containing twenty-nine variants, which were independently co-transfected with the canonical gRNA_4 in *p53^tdTomato^* mESC. Most mutations were neutral or negatively impaired editing efficiency. However, two variants, S53R and E401R, showed significantly elevated KO levels (23.8 ± 1.9% and 32.7 ± 0.8% vs 18.6 ± 1.9% control), with E410R increasing editing by 1.76-fold compared to the unmodified Cme2 protein (Figure 2G). The result was validated on a second target (gRNA_1), where the E410R mutation also enhanced Cme2 activity (Supplementary Figure 2F). To explore possible additive effects, the two mutations were combined to obtain a S53/E410R double-variant. Surprisingly, the double mutant exhibited a negative impact on editing compared to E410R alone, implying a functional interaction between S53R and E401R (Supplementary Figure 2F). However, these data demonstrate that E410R significantly improves Cme2 functionality.

We next sought to optimize nuclear localization sequence (NLS) composition. There is currently no consensus on the optimal NLS pattern for Cas proteins, as it varies by CRISPR system and application among other factors, though the addition of multiple NLSs has been shown to enhance editing in some cases [44–46]. In the present study, we designed four distinct arrangements each comprising a different number and type of NLSs (Supplementary Figure 2G). Among the combinations, Cme2_d04 design (3x c-myc NLS-Cme2-2xSV40 NLS) exhibited the highest editing activity in both the sites tested compared to the original NLS pattern (52.8 ± 2.3% vs 48.9 ± 3% with gRNA_1 and 29.2 ± 1% vs 18 ± 1% with gRNA_4). (Figure 2F). We therefore integrated the optimal NLS design with the E410R mutation to obtain the final optimized protein variant, termed Cme2_fd1.

Finally, we aimed to assess whether the improvements in Cme2 activity achieved through protein and sgRNA engineering were cumulative by combining Cme2_fd1 and Cme2_en_sg1. This iteratively engineered combination was named *en*Cme2 (i.e., enhanced Cme2). We compared *en*Cme2 with Cme2 using the twelve sgRNAs targeting the *tdTomato* gene employed in the initial Cme2 validation shown in Figure 1F. The data revealed a synergistic effect, leading to a significant improvement in editing efficiency for gRNA_1, gRNA_4 and gRNA11 with increases of 1.6-fold, 3.8-fold and 5.2-fold, respectively (Figure 2H). Surprisingly, we observed that gRNA_6, which originally exhibited only background-level activity, achieved a KO efficiency of up to 16.6% with *en*Cme2 (57.7-fold improvement over the control). Ultimately, we observed an average overall improvement in editing activities of 2.23-fold and 5.19-fold in the functional and non-functional sgRNA groups, respectively (Supplementary Figure 2H). *en*Cme2 therefore represents a rationally evolved compact Cas9 protein, with markedly increased ON-target activity compared to Cme2.

### ON-target validation and OFF-targeting assessment of *en*Cme2

To enable the facile optimization of Cme2 towards enhanced *en*Cme2, we previously used the expression of a reporter as a proxy for upstream indel generation, which underestimates activity. To investigate the actual ON- and OFF-target genomic editing, we first targeted multiple regions in the endogenous *p53* promoter and quantified indels using TIDE [47]. Using ten randomly designed sgRNAs, we observed *en*Cme2 exhibits superior activity compared to Cme2. Specifically, indels were detected in 7/10 targets, with a significant improvement in activity observed in six of these seven cases (Figure 3A). On average, *en*Cme2 exhibited 1.58-fold higher efficiency than unmodified Cme2 at the *p53* endogenous locus, confirming the advantageous impact of our engineering efforts.

**Figure 3.**
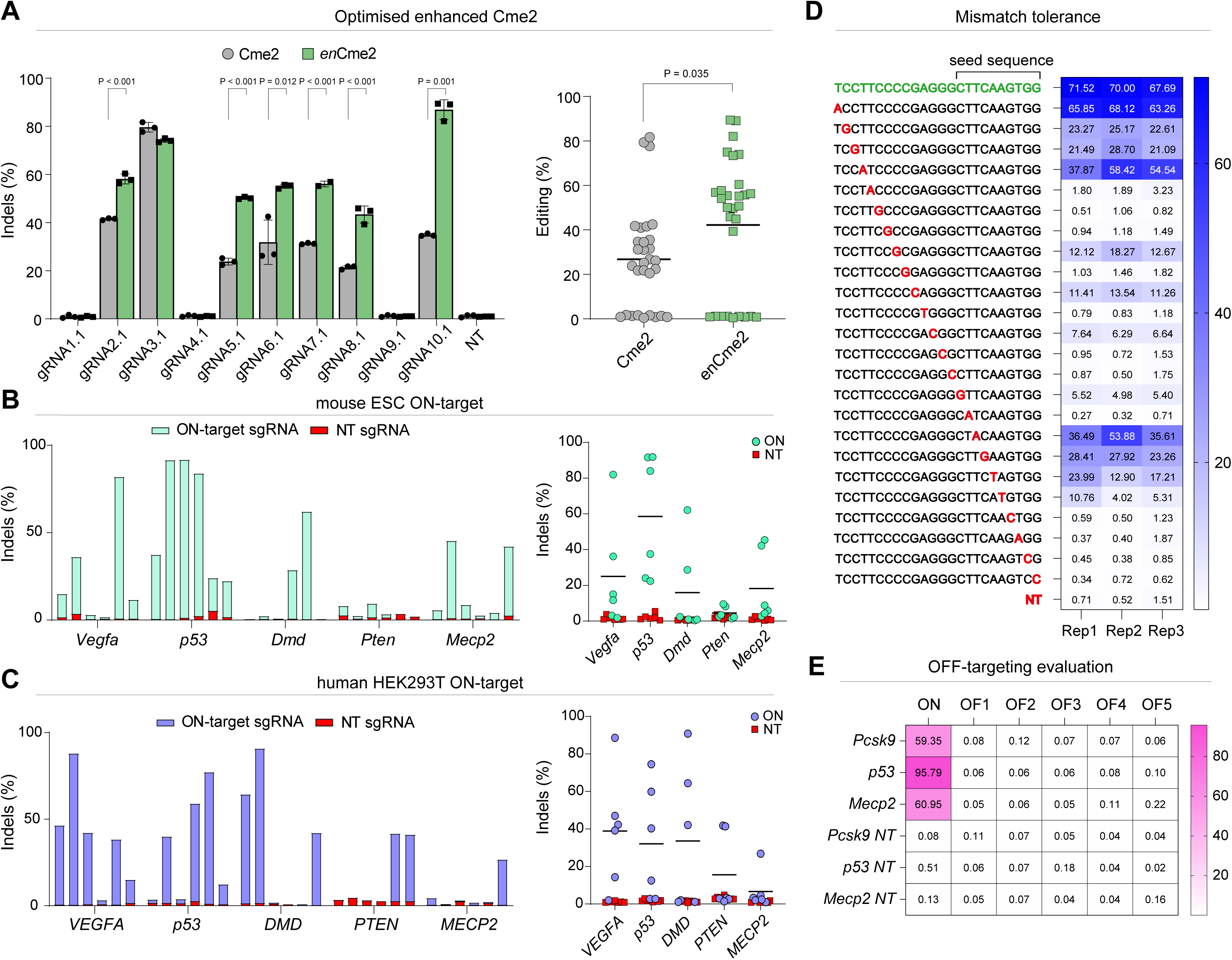
Evaluation of ON and OFF-target activity of *en*Cme2. (A) Histogram comparing the editing efficiency of *en*Cme2 and Cme2 at ten target sites within the endogenous *p53* gene in mESC. (n=3) (left). Average editing efficiency at the *p53* locus for *en*Cme2 and Cme2 (right). Two-group comparisons were performed by unpaired *t*-test. Error bars ± SD. (B-C) Quantification of Indels generated by *en*Cme2 in five endogenous loci in mESC (B) and HEK293T cells (C). Left panels: overlaid histograms showing editing using six distinct sgRNAs for each gene (*Vegfa, p53, Dmd, Pten, Mecp2*) in both cell lines. Right panels: average editing across the five different loci (six sgRNAs per gene). Controls are shown in red, bars represent mean values of ON-target activity in each gene. (D) Tolerance of *en*Cme2 to single-mismatched bases in gRNA_1 spacer. Fully matched ON-target spacer is indicated in green, mismatched bases are highlighted in red. Three biological replicates are shown individually. (E) Heat map showing the percentage of indels generated by *en*Cme2 at predicted off-target sites in mESC, relative to controls, as determined by targeted deep sequencing.

We next investigated *en*Cme2 activity on a set of therapeutically relevant targets in different cell types and species. We selected *Vegfa, p53, Dmd, Pten* and *Mecp2* and designed six sgRNAs for each gene based strictly on PAM sequence. We detected indel formation (normalized editing > 3%) in eighteen of the thirty total sites (60%) assayed in mESC, up to a total maximum efficiency of 91.8%. Average editing efficiency varied among genes, ranging from 4.6% ± 3.4% for *Pten* to 58.6% ± 34% for *p53*, with a total average editing of 24.4% ± 29.8% in mESC (Figure 3B). We then sought to validate the results in human cells, maintaining the same target genes. We designed an additional thirty human-specific sgRNA (six per gene) for validation in HEK293T cells. We observed indels in fifteen out of thirty sites (50%), with an overall average editing efficiency across all genes of 25.51% ± 28.49%, and the highest average activity observed in the *VEGFA* gene at 38.9% ± 29.3% (Figure 3C). Overall, *en*Cme2 demonstrated activity in more than 50% of the tested targets across both cell types, highlighting its potential application in research and therapeutically relevant contexts and across different species.

From a clinical perspective, editing efficiency is just one factor to consider when selecting a CRISPR system for a particular application. Among other factors, specificity is one of the most critical [48–51]. To evaluate this, we tested the effect of a single mismatch in the protospacer sequence on *en*Cme2 activity. We systematically introduced a single mismatch at each position of the gRNA_1 spacer and observed that, for the *tdTomato* gene, a single mismatch at positions 6, 7, 9, 11, 13, 14, 16, and 20-24 completely abolished editing activity. Similar to other CRISPR systems, *en*Cme2 tolerated mismatches in the PAM-distal region (positions 1-4) [49] [35] [19]; however, we noticed that mismatches in two positions (17-18) of the seed sequence were also tolerated, though they still negatively impacted activity (Figure 3D).

To assess off-target activity at endogenous genomic sites, we performed targeted deep sequencing on potential OFF-target sites identified using Cas-OFFinder [52]. As ON-target sites, we selected validated sgRNAs targeting *Pcsk9*, *p53*, and *Mecp2* genes. Notably, due to the restrictive PAM requirement of *en*Cme2, even the closest potential off-targets showed a high number of mismatches compared to the ON-target protospacer sequences. Indeed, the nearest OFF-targets for the *Pcsk9, p53,* and *Mecp2* spacers differed by at least 5, 6, and 6 nucleotides, respectively, from the intended target sites (Supplementary Figure 3). As expected, targeted deep sequencing analysis revealed no detectable indels at any of the 15 potential OFF-target sites analyzed, with editing levels comparable to the negative controls (p>0.05) (Figure 3E). These data suggest that compact *en*Cme2 exhibits exquisite ON-target specificity.

### Cme2 as an epigenome editing tool

As well as genome sequence editing, compact Cas9 proteins hold great promise for epigenome editing, whereby transcription levels are precisely modulated. We therefore explored the potential of utilizing baseline Cme2 as an epigenome editing tool. We first generated two Cme2 nickases (nCme2 D4A, nCme2 H576A) as well as a catalytically inactive “dead” Cme2 variant (dCme2 D4/H576A), which were fused with an optimized array of GCN4 repeats [23]. Our recent studies have shown this can recruit up to five copies of a catalytic domain of a chromatin modifier, or a repressive KRAB domain (iCRUSH), to amplify ON-target epigenome editing. Western blot analysis confirmed similar expression levels of the different nickase or dead *Cme2* variants (Supplementary Figure 1D). To test functional activity, we introduced the nCme2 systems together with the G9a catalytic core (G9a^scFV^) into p53^tdTomato^ mESC [53]. G9a^scFV^ programs the histone modification H3K9me2, which triggers promoter repression. Indeed, we observed a reduction of single-cell expression level when targeting two *p53* promoter sites with nCme2 D4A, but not with nCme2 H576A (Supplementary Figure 1E-F). Although modest, the observed activity highlights the potential use of Cme2 for epigenetic editing. Future work to iteratively evolve an enhanced *d*Cme2 (*en-d*Cme2) with optimal activity, analogous to the *en*Cme2 here, is therefore warranted.

### Genome editing in mouse embryos through single AAV delivery of enCme2

An inherent advantage of the compact size of *en*Cme2 is that the entire system can be packaged within a single AAV vector, making it particularly appealing for *in vivo* applications. As an exemplar use case of this advantage, we aimed to evaluate whether *en*Cme2 can generate transgenic mice through *ex vivo* exposure to an all-in-one rAAV vector carrying the entire CRISPR system (Figure 4A). To this end, we constructed two plasmids encoding *en*Cme2 and two different sgRNAs targeting *tdTomato* (gRNA_1 and gRNA_4), under the control of the EF1α core promoter and the hU6 promoter, respectively, resulting in a total size of approximately 4.5 Kb (p_hU6::gRNA(1)_EF1a::*en*Cme2; p_hU6::gRNA(4)_EF1a::*en*Cme2). These plasmids were packaged into rAAV1 particles (rAAV1:*en*Cme2-gRNA1, rAAV1:*en*Cme2-gRNA4) and optimized via transducing p53^tdTomato^ mESC at increasing concentrations. We quantified the KO efficiency via flow cytometry after 4 days (Figure 4B) and observed that gRNA_1, but not gRNA_4, exhibited modest activity in a concentration-dependent manner, with a maximum editing efficiency of 1.97% ± 0.83%. The low level of editing observed can be explained by the lack of cell selection and the particularly low transduction efficiency of mESC by AAVs, in line with the literature [54]. This was confirmed by the analysis of p53^tdTomato^ mESC transduced with rAAV1:Cre-eGFP vectors at the same doses used for the previous experiment, where we noticed a maximum of 1.09% ± 0.08% GFP-positive cells in the higher concentration group (Supplementary Figure 4A). Nevertheless, these data confirm that *en*Cme2/sgRNA packaged into a single rAAV are functionally competent to induce specific genome edits.

**Figure 4.**
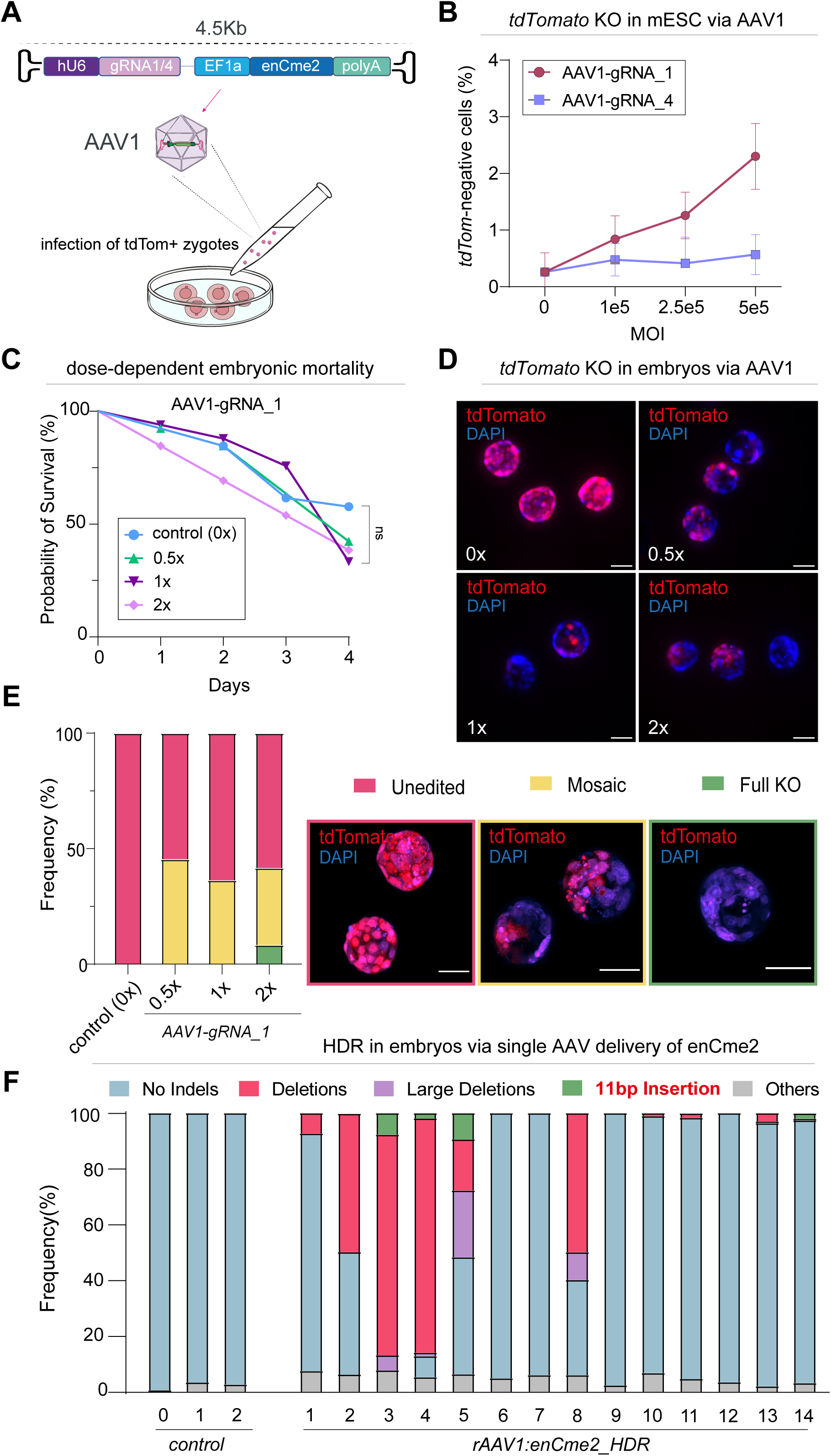
Single-AAV delivery of *en*Cme2 for efficient *in vivo* editing. (A) Schematic representing the strategy to perform *tdTomato* KO in both zygotes and mESC through rAAV1:*en*Cme2 delivery. (B) Assessment of editing at the *tdTomato* locus in mESC using increasing concentrations of rAAV1:*en*Cme2-gRNA1 (rAAV1-gRNA_1) and rAAV1:*en*Cme2-gRNA4 (rAAV1-gRNA_4). Each dot represents the mean ± SD of three independent biological replicates. MOI, multiplicity of infection. (C) Survival curve showing mortality levels of B6.Cg-Gt(ROSA)26Sor*^CAG-tdTomato^*embryos treated with increasing doses of rAAV1-gRNA_1 compared to untransduced zygotes. Ns indicates P-value > 0.05 by Mantel-Cox test. (D) Representative images of B6.Cg-Gt(ROSA)26Sor*^CAG-tdTomato^* embryos transduced with different rAAV1-gRNA_1 doses. DAPI in blue. Scale bar 50 μm. (E) Left: stacked histogram showing the prevalence of edited B6.Cg-Gt(ROSA)26Sor*^CAG-tdTomato^* embryos among four rAAV-dosage conditions, grouped according to the presence of tdTomato-negative cells in the embryos. Right: representative confocal images of the different groups. Scale bar 50 μm. (F) Stacked histogram displaying the percentage of different indel types in untreated or rAAV1:*en*Cme2_HDR-transduced C57BL/6J embryos. Large deletions: > 10 bp; Deletions: 1-10 bp; Others: reads with P-value > 0.05 or not analyzed due to insufficient alignment score. In panels C-D-E, 0.5, 1 and 2x concentrations indicate 6.58 × 10^9^, 1.31 × 10^10^, 2.62 × 10^10^ VG doses, respectively. 0x indicates the no-virus control. In panels D and E, images were acquired with a Leica Thunder and Nikon AX/AR microscopes, respectively.

We therefore employed rAAV1-gRNA_1 to transduce *in vivo* zygotes carrying a reporter cassette (B6.Cg-Gt(ROSA)26Sor^CAG-*tdTomato*)^) at different concentrations (6.58 × 10^9^, 1.31 × 10^10^, 2.62 × 10^10^ VGs per culture droplet). We observed no significant difference in the survival rate of embryos treated with the control (P > 0.05) regardless of the rAAV1 dose used, confirming the safety profile of rAAV vectors [10] (Figure 4C). To assess editing efficiency, embryos were examined at the blastocyst stage using confocal and epifluorescence microscopy. We noted the presence of partially or entirely tdTomato-negative embryos at all tested concentrations (Figure 4D and Supplementary Figure 4C), with the latter implying an early genomic edit (zygote or 2-cell stage) and thus timely *en*Cme2 action. Because tdTomato is a tandem-dimer composed of two copies of the tdTomato fluorophore [55], each sgRNA can induce two distinct cuts, which can lead to various gene rearrangements, complicating the generation of clear PCR products for sequencing (Supplementary Figure 4B). For this reason, we assessed the efficiency of *en*Cme2 based on the proportion of fluorescently-negative embryos relative to the total transduced. Embryos were classified into three groups: unedited, mosaic, or full KO. In the 6.58 × 10^9^ VGs, 1.31 × 10^10^ VGs, and 2.62 × 10^10^ VGs groups, the percentages of edited embryos were 45.4%, 36.4%, and 41.2%, respectively, with complete penetrance of KO across all embryonic cells observed in the highest concentration (Figure 4E). These data confirm the feasibility of deploying *en*Cme2 to generate transgenic mouse embryos via delivery in a single all-in-one rAAV.

Taking into account that the compact size of *en*Cme2 leaves approximately 300 bp of available space when packaged into an rAAV, we exploited this residual capacity to include an HDR template to assess the feasibility of achieving small knock-ins in mouse embryos through single AAV delivery. As a proof of principle, we designed an 11bp stop cassette containing two premature termination codons flanked by minimal homology arms (100bp). This premature stop cassette was inserted into an *en*Cme2 encoding plasmid harboring a sgRNA targeting exon 5 of the mouse *p53* gene (p_hU6::m_gRNA4-p53_EF1a::*en*Cme2_HDR), which previously showed high activity in mESC (Figure 3B). A packaged AAV1 (rAAV1:*en*Cme2_HDR) was co-incubated with C57BL/6J mouse zygotes (5 × 10^9^ VGs per culture droplet), with the exposed embryos cultured until the blastocyst stage, where indels were quantified via Sanger sequencing. To enable a precise classification of indel types and the expected 11 bp insertion, we categorized the indels into distinct groups (Figure 4F). We identified the presence of indels in 10 of the 14 tested embryos, with a total prevalence of 71.4% and a maximum efficiency of 100%. We observed variability in editing efficiency among different embryos, with small deletions preferentially occurring. However, importantly, the only insertion event detected was an 11 bp insertion. This perfectly matched the stop cassette sequence in 75% of cases, as confirmed by TIDER (pre-released TIDER code, see Methods). HDR occurred at a frequency spacing from 1.9% to 9.4% and an overall prevalence of 21.4%. The cumulative data support that the engineered variant *en*Cme2 is highly specific and can be used within a single-AAV delivery vector for *in vivo* genome editing and HDR knock-in.

## DISCUSSION

The use of compact CRISPR systems and their delivery via rAAV vectors for *in vivo* gene editing is emerging as a potentially transformative strategy for a wide range of clinical and research applications. Indeed, CRISPR-based therapy for sickle-cell disease has recently acquired FDA approval [56], while there is broad therapeutic use of rAAV-mediated delivery for remedial solutions, including for DMD [57] and Hemophilia A [58]. To facilitate combining the precision of CRISPR with the delivery potential of rAAV in the future, it is important to develop novel and compact CRISPR systems that expand the toolkit for distinct applications, as well as enhancing aspects such as targeting range and specificity. This will, in turn, amplify the therapeutic and research application space for CRISPR.

In this study, we assessed the activity of a set of previously characterized Cas9 orthologs, aiming to develop novel CRISPR tools for *in vivo* applications. We initially evaluated Cas9 orthologs editing across different cell types, revealing an overall low activity among the systems tested. However, we decided to focus on one specific ortholog, Cme2, based on its relatively higher efficiency and potentially advantageous PAM sequence. Through rational design and iterative strategies, we optimized the structure of both the Cme2 protein and its sgRNA to develop a more robust and effective system. The enhanced sgRNA architecture was achieved through a combination of structural modifications, including extending the crRNA–tracrRNA hybridization region, shortening a stem-loop structure formed by the 3′-tracrRNA, and correcting a bulge in the crRNA-tracrRNA duplex. These designs were intended to enhance sgRNA stability and improve its interaction with Cme2 protein, collectively resulting in a significant increase in ON-target activity. In parallel, we aimed to strengthen the interaction between the Cme2 protein and target DNA, and to facilitate RNP formation, by employing a rational protein-engineering approach [35]. This led to the development of a high-activity variant of the Cme2 protein. By combining the improvements achieved through protein and sgRNA engineering with further NLS design optimization, we established a final <u>en</u>hanced system, termed *en*Cme2. Compared to canonical Cme2*, en*Cme2 exhibited a substantial improvement in editing frequencies across both endogenous and exogenous targets, and demonstrated activity on targets where Cme2 was non-functional. Thus, the optimization strategies we employed are effective and highly versatile and provide a framework for improving emerging CRISPR systems. In this respect, it would be interesting to probe whether *en*Cme2 is able to improve the poor epigenome editing activity observed with Cme2, which is potentially linked with insufficient occupancy time at target DNA.

Large-scale validation of *en*Cme2 across endogenous targets revealed its ability to effectively edit a broad range of sites in diverse cell types and species. Notably, *en*Cme2 extended PAM sequence (5′ NNAGNGC) enhances specificity by significantly reducing the likelihood of off-target effects. Indeed, we observed that even the most closely related potential off-targets typically exhibit multiple mismatches compared to the on-target sequence, with the presence of a single mismatch often sufficient to abolish *en*Cme2 activity entirely. This makes our *en*Cme2 system particularly well-suited for applications where minimizing off-target activity is a priority, such as gene therapy applications. Moreover, by leveraging the compact size of the *en*Cme2 system, it is possible to package the entire construct into a single rAAV vector, leaving additional space for regulatory elements, tissue-specific promoters, or small HDR templates (up to ∼300 bp). Indeed, we demonstrated that *en*Cme2 can be efficiently delivered into mouse zygotes through a single AAV1 vector, achieving robust editing activity without any further manipulation or specialized equipment. To our knowledge, this study is the first to achieve a precise genomic KI in mouse embryos using a single rAAV vector that carries both the complete CRISPR system and its HDR sequence. Traditional pipelines for engineering KI mice models generally integrate electroporation, viral transduction and/or microinjection to deliver the various CRISPR components separately. Our one-vector approach drastically simplifies the workflow and reduces both time and expense, thus establishing a scalable and economical platform with the potential to be applied across diverse species.

In conclusion, *en*Cme2 offers significant advantages, including its compact size and low off-target propensity, albeit with a restricted pool of targetable sites due to its PAM. These features make it a versatile option for a variety of *in vivo* applications, with potential implications for both basic research and therapeutic development.

## METHODS

### PLASMID CONSTRUCTION

To generate Cas9 expression systems, human codon-optimized *cco*, *cme2*, *gsp*, *msc*, *nsp*, *sdo*, *tmo* Cas9-encoding genes and sgRNA modules were designed based on a previous publication [22]. SpCas9 coding region and sgRNA cassette were amplified from pX330-gRNA (Addgene #158973). PCR reactions were performed using Q5 High-Fidelity DNA polymerase (New England Biolabs). Cas9 nuclease-encoding fragments were loaded on an agarose gel (0.7%), purified using a Gel and PCR Clean-up Kit (Macherey-Nagel) and cloned into the pPB_TRE3G::dCas9^GCN4^_*EF1a::*TetOn-Hygro [23] recipient plasmid under the control of the TRE3G promoter via “Gibson assembly” using NEBuilder HiFi DNA Assembly Master Mix (New England Biolabs). sgRNA expression plasmids were generated by inserting the Cas9 ortholog-specific sgRNA scaffold sequence into a modified U6::gRNA_EF1a::BFP-Puro vector [23] carrying silent mutations at BbsI sites, as described above. Oligonucleotides encoding the spacer sequences were annealed and ligated to BbsI sites using a T4 DNA Ligase v2 kit (Takara). All spacer sequences used in this work are listed in Supplementary Data. The nCme2 and dCme2 vectors were constructed by replacing the dCas9 coding sequence of pPB_TRE3G::dCas9^GCN4^_*EF1a::*TetOn-Hygro plasmid with synthesized Cme2-encoding DNA fragments carrying site-specific mutations (Azenta Life Sciences) using Gibson assembly. To generate p_hU6::gRNA(1,4)_EF1a::*en*Cme2 plasmids, Cme2_fd1 protein variant and Cme2_en_sg1 modified scaffold coding sequences were sequentially cloned into an AAV packaging vector under the Elongation Factor 1α (EF1α) promoter and hU6 promoters, respectively. p_hU6::m_gRNA4-p53_EF1a::*en*Cme2_HDR was obtained by replacing p_hU6::gRNA(1)_EF1a::*en*Cme2 spacer sequence with m_gRNA_4 (targeting *p53 –* see Supplementary Data) and inserting the HDR module, consisting of the TGATGAATTAT stop cassette flanked by 100 bp homology arms. All assembled plasmids used in this work were amplified in chemically competent *E. col* cells, purified using a NucleoSpin Plasmid kit (Macherey-Nagel) and sequenced by Sanger (Genewiz) and/or Whole-Plasmid sequencing (Eurofins Genomics) to confirm correct assembly.

### sgRNA and PROTEIN ENGINEERING

All sgRNA engineering steps were performed by cloning synthetic DNA fragments (IDT DNA Technologies) carrying specific mutations in the Cme2 scaffold coding region into the U6::gRNA_EF1a::BFP-Puro plasmid via Gibson assembly. Spacers were subsequently inserted as described above. Sequences encoding the engineered sgRNA scaffolds used in this study are listed in Supplementary Data.

For the generation of Cme2 protein mutants, the 29 single-point mutations were organized into five groups according to their relative positions within the Cme2 coding sequence, such that mutations located in close proximity were included in the same group (see Supplementary Data). p_TRE3G-wtCme2-hygroR plasmid was then linearized using either AflII restriction enzyme or sgRNA/SpCas9 RNP complexes targeted at specific locations designed to be at the closest proximity of the five different mutation clusters. For the *in vitro* plasmid digestion using Cas9 RNP, 500ng of custom-designed Alt-R™ CRISPR-Cas9 crRNA (IDT DNA Technologies) was annealed with 1 µg of tracrRNA (IDT DNA Technologies) in rCutSmart buffer (New England Biolabs) at 95 °C for 5 min and subsequently complexed with 500ng of Alt-R™ S.p. Cas9 Nuclease V3 (IDT DNA Technologies) at room temperature for 12 min. 5µg of template plasmid were added to the RNP complex and incubated for 3 hours at 37 °C. Following gel purification, synthetic DNA fragments (IDT DNA Technologies) or oligonucleotides (Eurofins Genomics) carrying desired mutations were inserted using Gibson assembly into the linearized Cas9-encoding plasmid. The different NLS combinations were obtained through PCR amplification of the Cme2 coding region and subsequently inserted into NheI/PmeI-linearized pPB_TRE3G::dCas9^GCN4^_*EF1a::*TetOn-Hygro plasmid using Gibson assembly.

### GENERATION OF *AAVS1^-tdTomato^*HEK293T REPORTER CELL LINE

The *AAVS1^-tdTomato^* HEK293T cell line was generated by introducing a *tdTomato* expression cassette into intron 1 of the AAVS1 gene through a CRISPR-based approach. The expression cassette, in which the *tdTomato* gene was driven by the EF1α promoter and followed by the Woodchuck Hepatitis Virus Posttranscriptional Regulatory Element (WPRE) and the Human Growth Hormone Polyadenylation signal (hGH polyA), was amplified via PCR from a donor vector (Addgene #67527) using appropriate ultramer DNA oligonucleotides carrying 100 bp-long overhangs complementary to the insertion site. 800 ng of the HDR template were nucleofected into 2×10^5^ HEK293T cells (ATCC, CRL-3126) along with 10 µg of Alt-R™ SpCas9 Nuclease pre-complexed with 5 µg of Alt-R™ CRISPR-Cas9 crRNA (spacer sequence: 5’ ACCCCACAGTGGGGCCACTA) and 5 µg of tracrRNA using a 4D-Nucleofector X Unit (Lonza Bioscience) following manufacturer’s instructions. Following electroporation, cells were routinely cultured for 4 weeks and subsequently FACS sorted to isolate single tdTomato-positive cells. After 3 weeks of expansion, candidate lines were screened for correct monoallelic integration through junction PCR and Sanger sequencing (Genewiz).

### CELL CULTURE AND TRANSFECTION

Heterozygous *p53^-tdTomato^* murine embryonic stem cells, obtained from a previous publication [23], were routinely cultured on gelatin-coated plates (0.1% gelatin, Sigma) and maintained in titrated t2i/L media prepared by supplementing NDiff 227 medium (Takara, Y40002) with 1 μM PD0325901 (Axon Medchem), 3 μM CHIR99021 (Axon Medchem), 1,000 U/mL leukemia inhibitory factor (EMBL PEPC facility), 1% FBS (Sigma), and 1% penicillin-streptomycin (Gibco). *AAVS1^-tdTomato^* HEK293T cells were cultured in Dulbecco’s modified Eagle’s medium (DMEM, Gibco) supplemented with 10% FBS, 1% penicillin-streptomycin solution and 1% L-Glutamine (Gibco). Cells were maintained at 37 °C in a humidified atmosphere with 5% CO_2_ and were passaged every 2/3 days by dissociation with TrypLE (Thermo Fisher Scientific, 12604013). Media was changed daily. Cells were transfected using Lipofectamine 3000 (Invitrogen) following the manufacturer’s instructions. Briefly, 0.8 µL of Lipofectamine 3000 (Invitrogen) was mixed with 350 ng of Cas9, 25 ng of PiggyBac transposase, and 50 ng of sgRNA-encoding plasmid(s). The mixture was incubated for 15 min and applied to cells at ∼70% confluency in 48-well culture plates. For the epigenome editing experiments, 100 ng of G9a-expressing plasmid was added to the mix. Transfected cells were subsequently selected with hygromycin B (200 μg/mL) for 5 days and puromycin (1.2 μg/mL) for 3 days. Neomycin (300 1.2 μg/mL) was added for 3 more days in G9a-transfected cells. Cas9 expression was then induced for 3 days using doxycycline (DOX) (100 ng/mL) before either flow cytometry or genomic DNA extraction.

### FLOW CYTOMETRY

To quantify knockout efficiency in tdTomato-expressing cell lines, cells were first detached using TrypLE and resuspended either in FACS buffer (10% FBS in PBS). Analysis was performed using an Attune NxT Flow Cytometer (Thermo Fisher Scientific) and data were analyzed with FlowJo v10.5.3 (Tree Star, Inc.). Knockout efficiency was calculated based on the percentage of tdTomato-negative cells over the total. Cell sorting was performed using a FACS Aria III (Becton Dickinson) flow cytometer.

### INDEL QUANTIFICATION

Genomic DNA was extracted from cells using a Quick-DNA Microprep Plus kit (Zymo Research) following the manufacturer’s instructions. Genomic regions of interest were amplified using target-specific primers (Eurofins Genomics) with either Q5 or primeSTAR GXL DNA polymerase (Takara Bio). Amplicons were purified from agarose gel and Sanger sequenced. Indel analysis was performed using TIDE, as previously described [47]. For the validation of *en*Cme2 at endogenous loci in mESC and HEK293T cells, normalized editing was calculated as the indel frequency of the ON-target sample minus that of the matched control. For targeted deep sequencing, gRNA-targeted regions were amplified with GXL polymerase as described above. Purified amplicons were subsequently labeled with P5+P7 Illumina adapters through a second PCR round and sequenced on an Illumina MiSeq i100 System (316-bp paired-ends reads) (Genomics Core Facility, EMBL). Indels were quantified within a 10 bp window from the cut site using CRISPResso2, as discussed elsewhere [59].

### WESTERN BLOT

Protein extraction was performed by resuspending the cell pellet in RIPA buffer (150 mM NaCl, 1% NP-40, 0,5% sodium deoxycholate, 0,1% SDS, 50mM Tris, SIGMA) containing Protease/Phosphatase inhibitors (Roche) and Pierce Universal Nucleases (Thermo-Scientific), incubated at 37 °C for 30 min and subsequently centrifuged. Cell lysis supernatant was collected, and the protein concentration was measured using a Pierce BSA protein assay kit (Thermo-Scientific). 15 μg of protein extract was mixed with 7,5 μL of Bolt LDS Sample Buffer (Novex) and 3 μL of NuPAGE reducing agent (Invitrogen), incubated at 95 °C for 5 min and loaded on a 4-12% Tri-Glycine Plus WedgeWell Gel (Invitrogen). Electrophoresis separation was carried on for 1 hour at 90V and subsequently for 1 hour at 110V in Tris-Gly SDS Running Buffer (Novex). Proteins were transferred to a PDVF membrane (ThermoFisher) using an iBlot3 stack (Invitrogen) and incubated in 5% milk/PBS-T (0,05% Tween 20) for 1 hour at room temperature. FLAG-tagged proteins were detected using an ANTI-FLAG M2 IgG1 antibody (Sigma, F3165) diluted 1:1000 in 0,5% milk/PBS-T, while β-actin and β-tubulin, used as internal loading controls, were probed with an anti-β-actin (Sigma, A5441) and an anti-β-tubulin (Sigma, T4026) diluted 1:2000 in 0,5% milk/PBS-T, respectively. After overnight incubation at 4 °C with primary antibodies, membranes were washed 3 times in PBS-T and incubated with peroxidase-conjugated secondary antibody (Jackson ImmunoResearch) diluted 1:10,000 in PBS-T for 1 hour at room temperature. After 3 washes in PBS-T, membranes were treated with Pierce ECL Plus WB Substrate (ThermoFisher) for 5 min and imaged with AMERSHAM ImageQuant 800.

### RECOMBINANT AAV VECTORS

rAAV1:*en*Cme2-gRNA1, rAAV1:*en*Cme2-gRNA4 and rAAV1:*en*Cme2_HDR vectors were produced and purified by VectorBuilder GmbH (Neu-Isenburg). rAAV1:Cre-eGFP vector, carrying a CRE-eGFP expression cassette driven by the CB6 promoter, was kindly provided by the Gene Editing and Virus Facility, EMBL Rome. All procedures involving the use of rAAV vectors containing complete CRISPR expression systems (nuclease + sgRNA expression modules) were conducted in a BSL-2 environment, in accordance with EMBL’s internal regulations.

### EMBRYOS and mES CELLS TRANSDUCTION

About 2×10^4^ p53^tdTomato^ mESC were plated in gelatin-coated 48-well cell culture plates and immediately infected with either rAAV1:*en*Cme2-gRNA_1, rAAV1:*en*Cme2-gRNA_4 or rAAV1:Cre-eGFP vectors at multiplicities of infection (MOI) of 10^5^, 2.5×10^5^ and 5×10^5^ viral genomes (VGs) per cell. Media was changed 12 hours after transduction and, subsequently, daily. Fluorescence analysis was performed 4 days post-infection via flow cytometry.

The B6.Cg-Gt(ROSA)26Sor^CAG-*tdTomato*^ reporter mice line was generated by crossing the B6.Cg-Gt(ROSA)26Sor^tm14(CAG-tdTomato)Hze/^J reporter line (Jackson Laboratory, #007914) with a ubiquitously expressed Cre-driver line to excise the loxP-flanked stop cassette and achieve constitutive expression of tdTomato. All mice were maintained on a C57BL/6J genetic background to preserve genetic consistency. For viral transduction, B6.Cg-Gt(ROSA)26Sor^CAG-*tdTomato*^ or wild-type C57BL/6J zygotes were thawed and equilibrated in EmbryoMax KSOM Advanced medium (Sigma) overlayed with mineral oil (Life Global) for 1 hour at 37 °C in a tissue culture incubator containing 5% CO_2_. Embryos were then transferred in 25 μl drops of KSOM under mineral oil and incubated for 6 hours with 6.58×10^9^, 1.31×10^10^ and 2.62×10^10^ VGs of rAAV1:*en*Cme2-gRNA_1 or 5×10^9^ VGs of rAAV1: *en*Cme2_HDR, previously pre-mixed with 25 μl of KSOM. Subsequently, embryos were washed 5 times with EmbryoMax M2 (Sigma) and transferred into 500µL KSOM drops in 60mm center well culture dishes (Corning). Zygotes were cultured for 4 days to reach the blastocyst stage and subsequently analyzed. Embryo mortality was assessed basis, and any non-viable embryos were discarded from the culture. rAAV transduction was performed in a BSL-2 facility until completion of the wash steps.

### EMBRYOS IMAGING and ANALYSIS

The efficiency of *tdTomato* KO in embryos was qualitatively assessed through fluorescence analysis relative to untransduced controls using epifluorescence microscopy (Leica DMi8). Subsequently, representative embryos from each experimental group were selected for DAPI staining. Briefly, embryos were washed in PBS, fixed with 4% PFA at 37 °C for 30min and incubated with DAPI solution (1 μg/mL) for 30 min. After two washes in PBS, embryos were imaged with a Nikon AX/AR scanning confocal microscope and a Leica Thunder Epifluorescence microscope. Images were analyzed using ImageJ v1.54.

To extract genomic DNA, embryos were incubated at 56 °C for 10 min, followed by incubation at 95 °C for 10 min in a lysis solution consisting of 100 mM Tris-HCl (8.3 pH), 0.45% Tween-20, 0.1 mg/mL gelatin, 50 mM KCl, 2.5 mM MgCl_2_ and 100 μg/mL Proteinase K. The *tdTomato* gene was then amplified using GXL polymerase with the following primers: tdTom_fw: (5’) AGCAAGGGCGAGGAGGTCATC; tdTom_rv: (5’) CTTGTACAGCTCGTCCATGC. 100ng of PCR products were mixed with Midori Green Direct solution (Nippon Genetics), loaded on a 0.7% agarose gel and imaged (Gel Doc XR+, Bio Rad). The *p53* gene was amplified as described above using primers p53_fw: (5’) caatggtgcttggacaatg and p53_rev: (5’) gctgttaaagtagaccctgg. PCR products were gel-purified, sequenced through Sanger sequencing and indel prevalence analyzed via TIDE [47]. Correct HDR prevalence was confirmed using a pre-release TIDER code (kindly provided by Andres Marco, Bioinformatics Lead at Data Curators BV).

### STATISTICAL ANALYSIS

Two-tailed Student’s *t*-tests and Mantel-Cox tests were performed using GraphPad Prism v9.5.1. The meaning and values of error bars, horizontal lines, data points in box and dot plots, P-values, as well as the number of replicates, are specified in each accompanying figure legend. In this study, sample sizes were not determined by any statistical method and experiments were performed without blinding.

## DATA AVAILABILITY

All data supporting the findings of this study, including the sequences of the plasmids used, are available from the corresponding authors upon request. Next-generation sequencing data generated in this study have been deposited in the European Nucleotide Archive (ENA) under accession number XXXXXXXX.

## ACKNOWLEDGEMENTS

We thank Virginijus Siksnys (University of Vilnius), Giedrius Gasiunas, Tomas Urbaitis and Miglė Štitilytė (CasZyme, Vilnius) for their support in this project. We gratefully acknowledge the Gene Editing & Virus Facility (EMBL Rome), particularly Maj Simonsen Jackson and Michela Ascolani, for support with mouse embryo experiments, as well as the Light Imaging, Flow Cytometry (EMBL Rome), and Genomics Core (EMBL Heidelberg) Facilities. We are grateful to Vladimir Benes (EMBL Heidelberg) and Antony Adamson (University of Manchester) for their valuable advice and support. We thank Andres Marco, Bioinformatics Lead at Data Curators BV for providing pre-release TIDER code. The authors declare no conflicts of interest. This project has received funding from the European Union’s Horizon 2020 research and innovation programme under the Marie Skłodowska-Curie grant agreement No 945405.

---

**Supplementary Figure 1.**
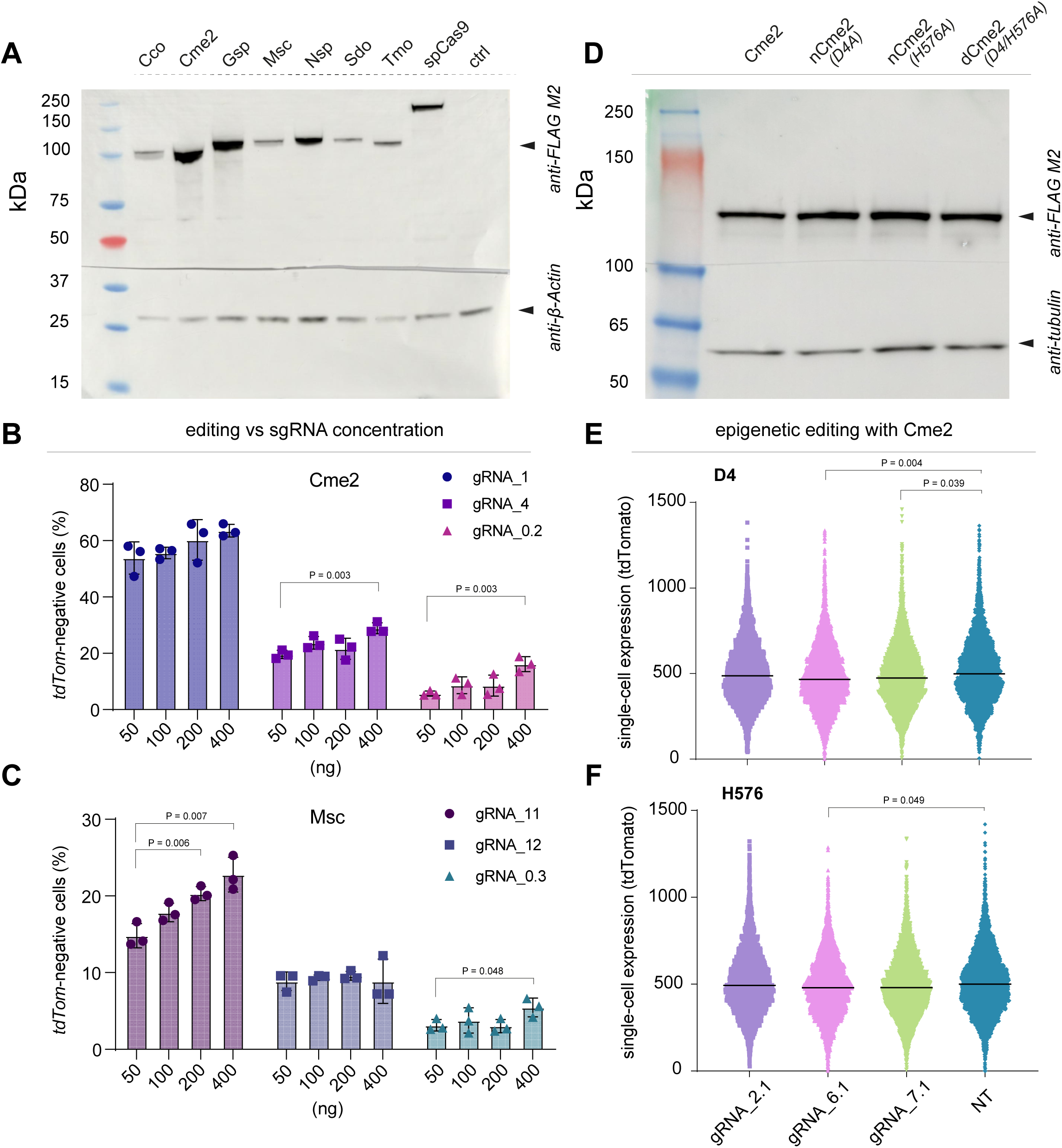
(A) Western blot analysis showing expression levels of different Cas9 orthologs in whole-cell lysates from p53*^tdTomato^* mESC transfected with the indicated Cas9-expressing plasmids. β-Actin served as a loading control. Ctrl represents untransfected cells. (B-C) Histograms showing Cme2 and Msc editing efficiencies in mESC transfected with increasing amounts of sgRNA-encoding plasmids. P-values were obtained using unpaired *t*-tests over three independent biological replicates; Error bars ± SD. (D) Western blot analysis showing *n*Cme2 and *d*Cme2 protein levels compared to Cme2. Cas9 proteins were fused with a 5x CGN4s repeats array, expressed in mESC and extracted 3 days after DOX induction. Tubulin was probed as a normalizing control. (E-F) Single cell expression of tdTomato after G9a-mediated (G9a-CDscFV) H3K9me2 using three sgRNAs targeting *p53* promoter or NT, employing Cme2 D4A (E) and H576A (F) nickases as DNA binding domains. *n*Cme2(s) were fused with a 5x CGN4s motif array, which enables G9a-CDscFV recruiting. Each data point indicates a cell and for each column, 1500 cells from three independent biological replicates were pooled. Bars represent the median. Unpaired *t*-test was performed on the mean of the medians of the three replicates.

**Supplementary Figure 2.**
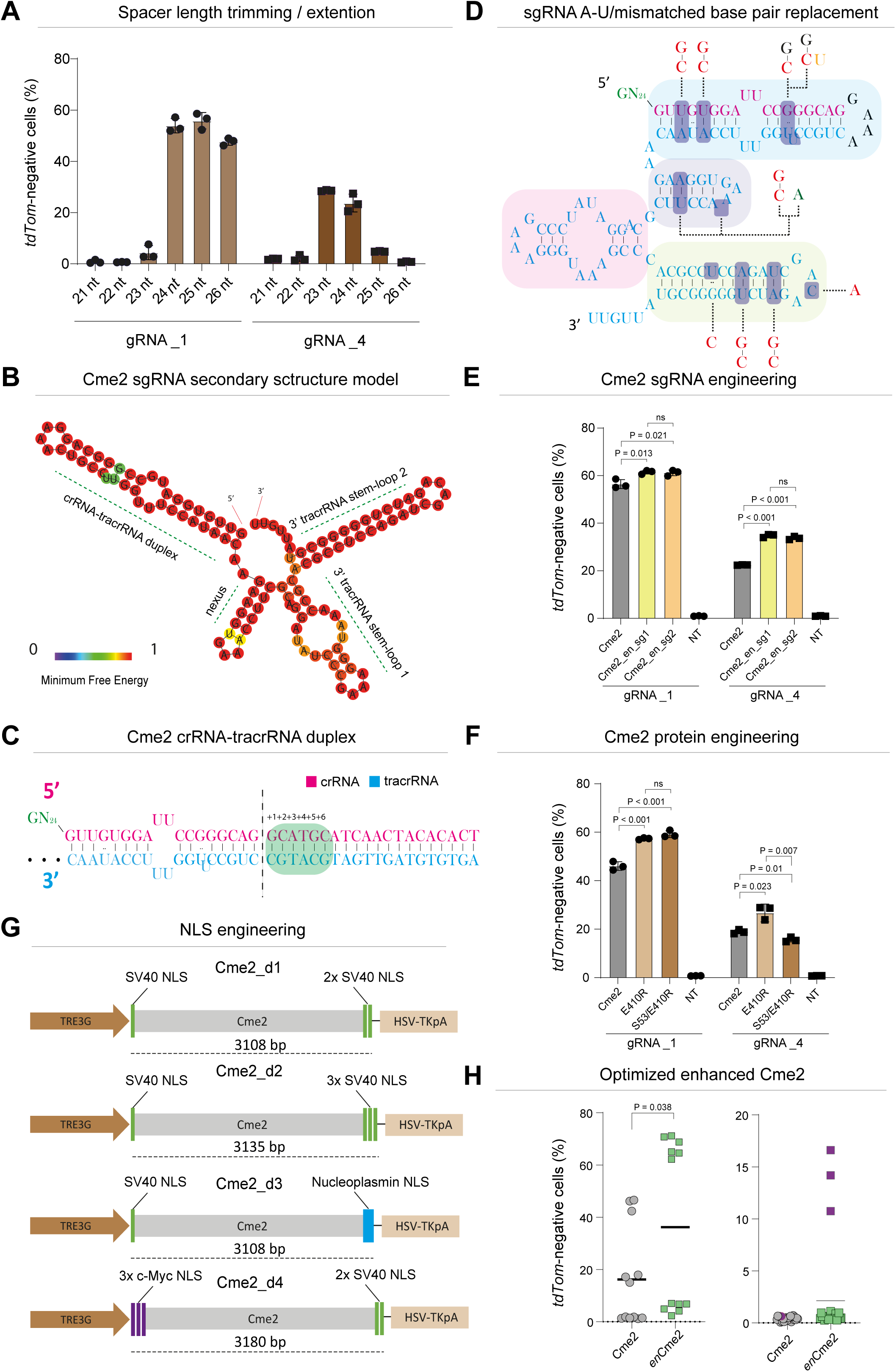
(A) Variation of Cme2 editing efficiency as a function of spacer length (21–26 nt) using gRNA_4 and gRNA_1. (B) Predicted minimum free energy (MFE) secondary structure of the Cme2 sgRNA scaffold. Model generated with the RNAfold web server. (C) Sequence of the native Cme2 crRNA–tracrRNA duplex. The vertical line marks the junction where the duplex was cleaved and rejoined via a GAAA tetraloop to generate the canonical sgRNA structure. Duplex extensions examined in this study (+1 to +6 bp) are highlighted in green. (D) Structure of the canonical Cme2 sgRNA, with main components enlightened as follows: blue (crRNA–tracrRNA duplex), purple (tracrRNA nexus), pink and yellow (3’ tracrRNA stem-loops 1 and 2, respectively). Ten modifications are indicated throughout the whole structure. Modified residues are color-coded: red for substitutions, green for insertions and orange for deletions. (E) Combinational effects of Cme2_a1 + Cme2_b10 + Cme2_c9 + either Cme2_b3 or Cme2_b4 sgRNA variants (to obtain Cme2_en_sg1 and Cme2_en_sg2, respectively) on Cme2 editing activity in mESC. P-values were obtained using unpaired *t*-tests over three independent biological replicates; Error bars ± SD. (F) Histogram showing the relative gene-editing efficiencies in mESC of the Cme2 E410R single-mutant variant and the Cme2 S53/E410R double-mutant variant. P-values were obtained using unpaired *t*-tests over three independent biological replicates; Error bars ± SD. (G) Schematic showing the various NLS configurations tested to improve Cme2 editing efficiency. Green bars indicate NLS sequences derived from the SV40 large T antigen, cyan bars represent NLS from *Xenopus* nucleoplasmin, and purple bars denote NLS from the human c-MYC protein. The canonical NLS configuration comprises two SV40-derived NLS motifs positioned at the N- and C-termini of the *cme2* coding region (H) Comparison of the average editing efficiencies of Cme2 and *en*Cme2 when guided by functional (left panel) or non-functional (right panel) sgRNAs directed to twelve independent *tdTomato* target sites. gRNA_6, which showed activity only with *en*Cme2, is indicated in purple. Two-group comparison was performed by unpaired *t*-test. Error bars ± SD.

**Supplementary Figure 3.**
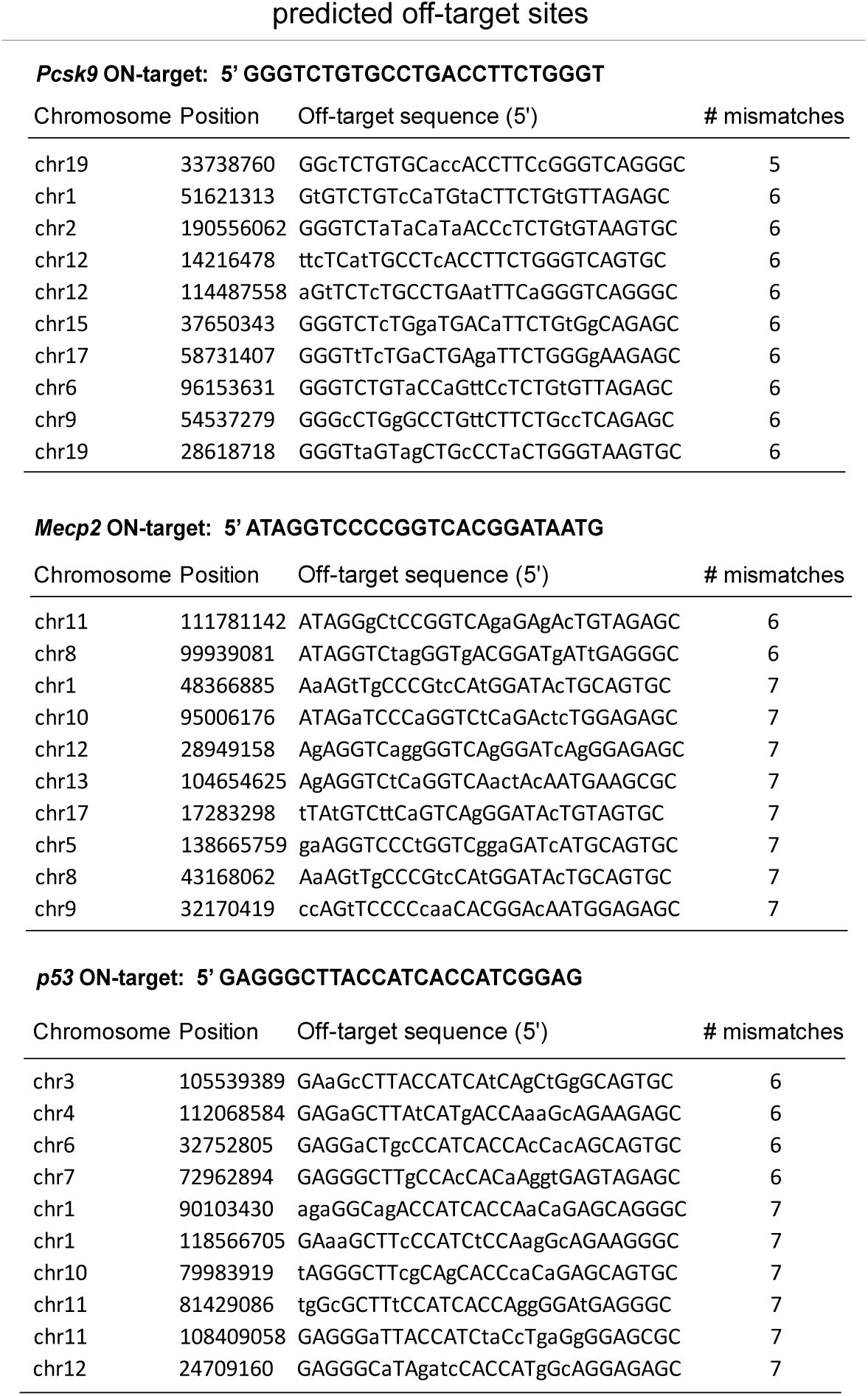
Tables listing the ON-target spacer sequences of *Pcsk9, p53*, and *Mecp2*, together with their predicted off-target sites identified using Cas-OFFinder and ranked according to the number of mismatches. Single-nucleotide mismatches are shown in lowercase.

**Supplementary Figure 4.**
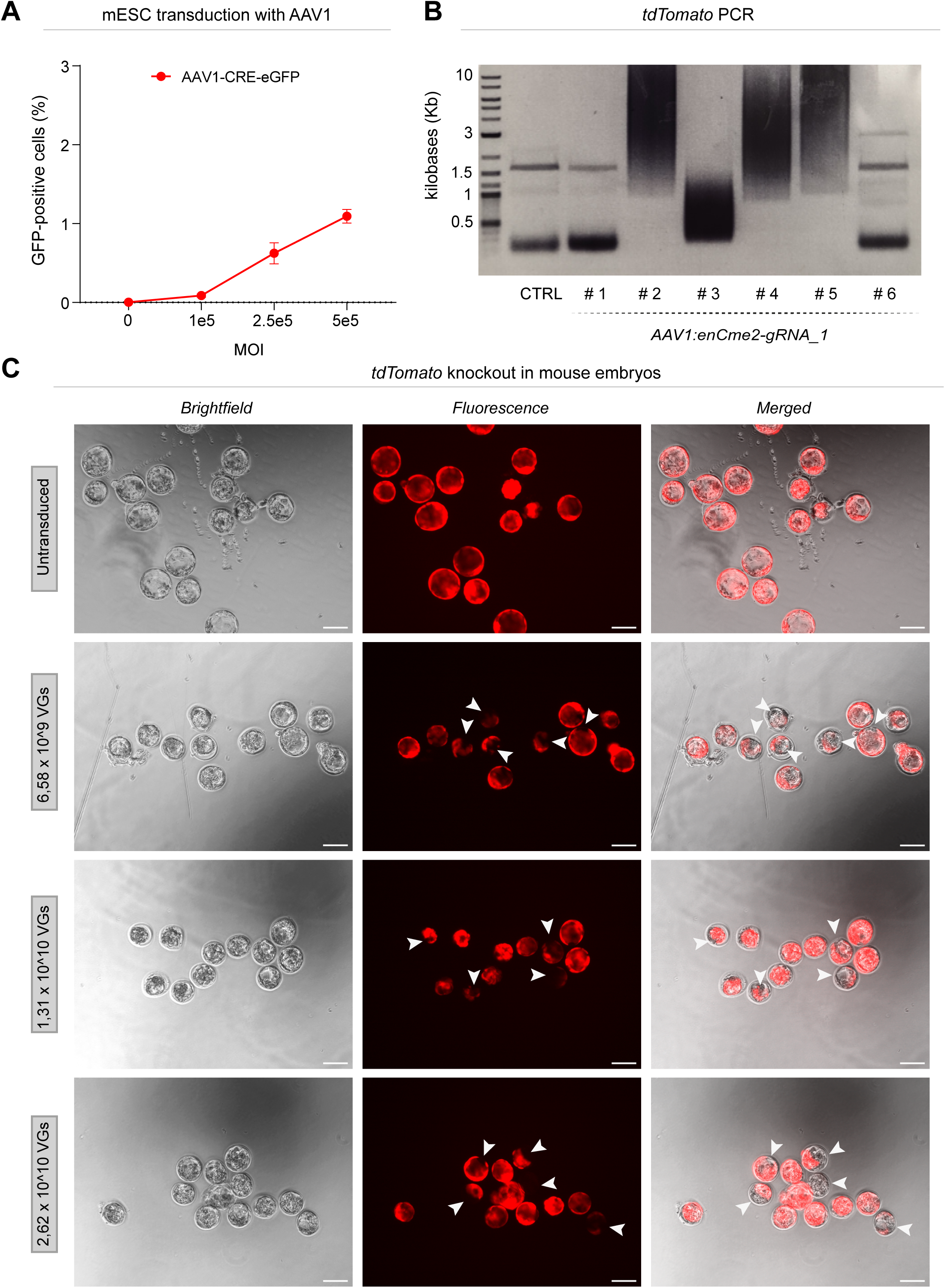
(A) Flow cytometric quantification of GFP-positive mESC four days after transduction with increasing doses of rAAV1-CRE-eGFP. (B) Agarose gel (0.7%) showing PCR amplicons generated from genomic DNA extracted from B6.Cg-Gt(ROSA)26Sor*^CAG-tdTomato^* embryos four days after transduction with 1.31 × 10¹⁰ VGs of rAAV1:*en*Cme2–gRNA_1 (samples 1–6) or from untreated controls (CTRL). The unedited *tdTomato* amplicon is approximately 1.5 Kb. (C) Epifluorescence images of B6.Cg-Gt(ROSA)26Sor*^CAG-tdTomato^* embryos transduced with increasing concentrations of rAAV1:*en*Cme2–gRNA_1, acquired four days post-transduction. White arrows indicate edited embryos. Scale bars, 100 μm.

